# XC-ID: De novo identification of the active X chromosome in single-cell RNA-seq

**DOI:** 10.64898/2025.12.05.692676

**Authors:** Jennifer Li Hui Jiang, Jesse Gillis

## Abstract

**Motivation:** X chromosome inactivation (XCI) is an epigenetic process that equalizes X-linked gene dosage between females (XX) and males (XY). During early development, one X chromosome in each cell is randomly silenced and clonally inherited, producing a stable mosaic of two epigenetically distinct cell lineages. This mosaicism provides a natural internal control for studying cell-intrinsic regulatory differences between X lineages. However, identifying the active X chromosome in single cells remains difficult due to sparse allelic coverage, dependence on pre-phased references, and biological variability from XCI escape and skew.

**Results:** We present **XC-ID** (**X C**hromosome inactivation **ID**entifier), a scalable computational framework for de novo identification of the active X chromosome from single-cell RNA-seq data. XC-ID employs a simulated-annealing algorithm to infer X-linked haplotype structure directly from allelic counts, followed by bootstrap-based confidence estimation to filter uncertain cell assignments. Applied to single-nucleus RNA-seq data from a female *Mus musculus* hybrid with known genotype, XC-ID achieved >99% accuracy in predicting the active X chromosome. The method remains robust to allelic noise, sequencing errors, and sparsity, and differential expression between inferred X lineages reveals biologically coherent dosage-compensation patterns.

**Availability:** XC-ID is available as a Python package with both API and command-line support at https://github.com/jlhjiang/XC-ID.

## Introduction

X chromosome inactivation (XCI) is an epigenetic mechanism that balances X-linked gene dosage between females (XX) and males (XY). Early in development, the transcription of one X chromosome in each cell is randomly silenced (Lyon 1961). This silenced state is clonally inherited through subsequent cell divisions, producing a mosaic of two epigenetically distinct cell lineages: one expressing the maternal X and one the paternal X (Lyon 1962).

This mosaicism provides a powerful internal control for studying allele-specific regulatory differences and cell-autonomous disease mechanisms. Such studies using bulk RNA-sequencing studies measure averaged allelic ratios but masks the cell-specific nature of XCI (Tukiainen *et al*. 2017; Werner *et al*. 2022; Werner, Hover and Gillis 2024). Single-cell RNA-seq (scRNA-seq) allows direct identification of the active X allele in individual cells through parental single-nucleotide polymorphisms (SNPs). Yet, current approaches are limited by dependency on pre-phased variants, complex custom pipelines, or computationally intensive models that scale poorly with large datasets.

Here, we introduce **XC-ID** (**X C**hromosome inactivation **ID**entifier), a lightweight framework for de novo phasing of the X chromosome and confident cell-level assignment of active X identity from scRNA-seq data. XC-ID employs simulated annealing to infer haplotypes (i.e. grouping SNPs by parental origin) without external phasing, followed by bootstrap resampling to filter uncertain cell assignments. This combination yields robust and accurate assignments even under high sparsity or poor sequencing quality.

The workflow (**Figure 1A**) includes constructing allele-specific SNP matrices from aligned BAM/VCF files, simulated-annealing phasing, bootstrap-based confidence estimation, and labeling each cell’s active X chromosome.

**Figure 1.**
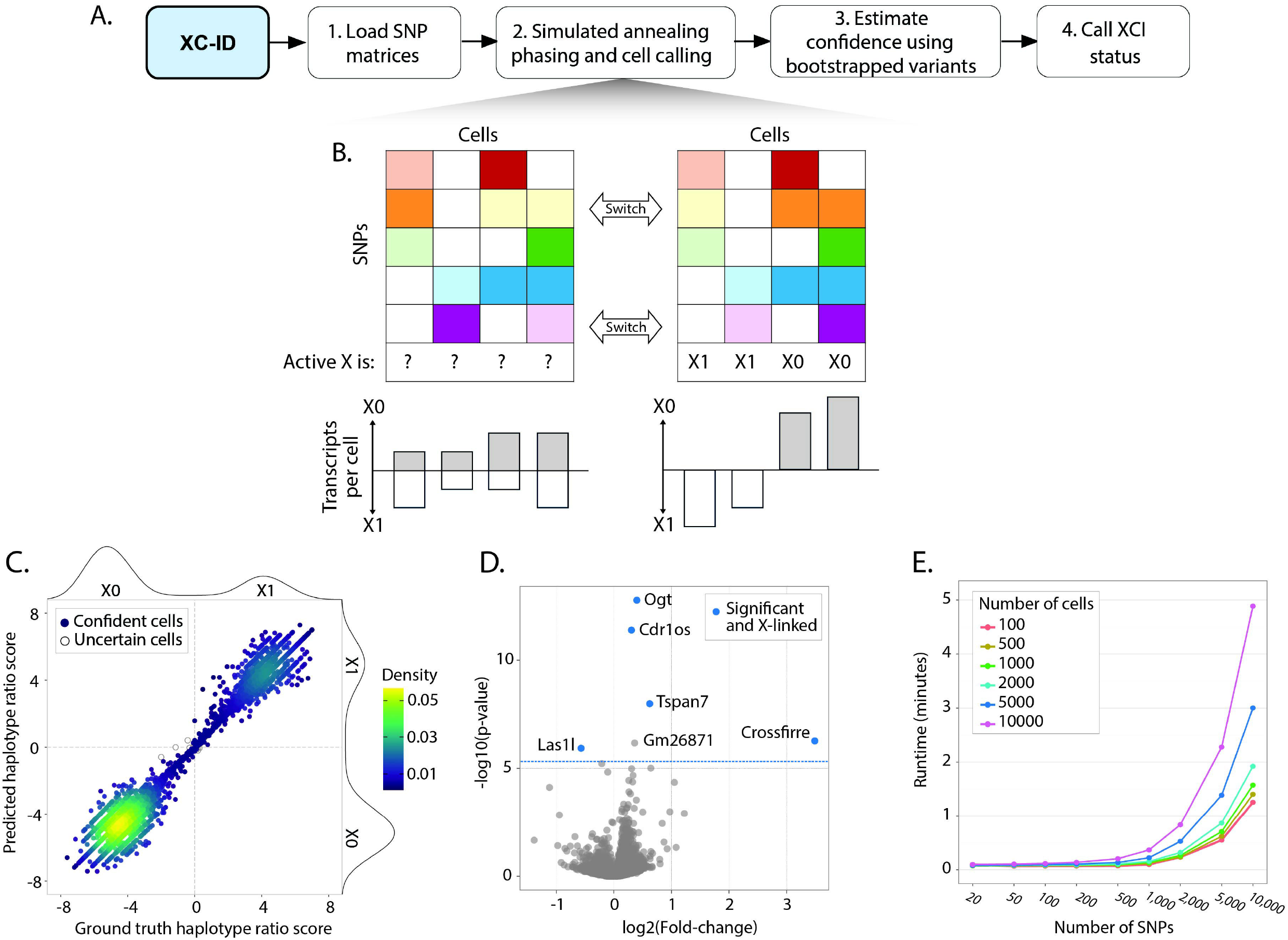
**(A)** Workflow of the XC-ID pipeline. SNP count matrices are loaded from STAR+WASP aligned BAM files, haplotypes are inferred using simulated annealing, confidence is estimated by bootstrapping, and per-cell X-inactivation status is assigned. **(B)** Simulated-annealing phasing schematic. The state of each SNP is iteratively “flipped” between the two candidate X haplotypes until transcript concordance across cells is maximized (i.e. each cell’s reads align predominantly to one haplotype). **(C)** Predicted versus ground-truth haplotype-ratio scores across all cells. Confidence labels were derived from bootstrap resampling. Pearson correlation = 0.99. **(D)** Volcano plot of differential expression between cells assigned to the two active-X lineages. X-linked and significant genes are highlighted. **(E)** Runtime performance of XC-ID across datasets varying in cell number and SNP count (log-scaled on the x-axis).

## Implementation

### Input and preprocessing

XC-ID accepts aligned scRNA-seq data and variant calls in standard formats. Reads are aligned using *STAR* with *WASP* filtering to remove allele-mapping bias (Dobin *et al*. 2013; van de Geijn *et al*. 2015; Asiimwe and Alexander 2024). We provide documentation for de novo SNP calling using *bcftools* (Li 2011), but variant information from databases such as 1000 Genomes or gnomAD can also be used. Informative bi-allelic SNPs are filtered and represented as two matrices

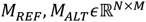

where *N* is the number of cells and *M* the number of SNPs. Each entry *M*_*REF*_(*i, m*) and *M*_*ALT*_(*i, m*) records the number of reads supporting the REF or ALT allele for cell *i* at SNP *m*.

### Simulated annealing haplotype inference

Phasing X variants is conceptually intuitive, as there are only two cellular states (maternal vs paternal X active). This binary structure can be leveraged to group heterozygous X-linked SNPs into two haplotypes. In practice, allele-specific analysis in scRNA-seq is often limited by transcriptomic sparsity, technical artifacts, XCI skew, and/or escape genes. To address this issue, XC-ID employs a simulated annealing algorithm to overcome local minima introduced by noise and error during haplotype inference.

XC-ID initializes a random binary phasing vector *v*_0_*∈*{*0,1*}^*M*^, representing the allele configuration across SNPs (**Figure 1B**). At each iteration *k*, a single SNP index *m* is randomly selected, its state flipped, and the configuration accepted according to the Metropolis-Hastings criterion with a geometric cooling schedule. The energy function maximizes allelic concordance across cells:

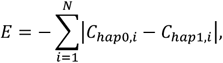

where

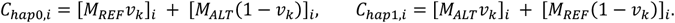

### Haplotype scoring

Following convergence of SNP groupings, per-cell haplotype preference is quantified as:

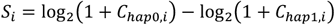

where negative and positive *S*_*i*_ values indicate preference for haplotype 0 or 1, respectively. The resulting bimodal distribution of *S*_*i*_ scores across cells delineates the two X chromosome haplotypes (**Figure 1C**).

### Bootstrap confidence estimation

XC-ID performs a bootstrap procedure to assess the per-cell confidence of X assignment. For each bootstrap, SNPs are resampled with replacement, the SA phasing is rerun, and a new set of per-cell scores is generated. Because the algorithm is direction-agnostic, each bootstrap score vector is realigned to the observed configuration based on SNP-level similarity weighted by transcript coverage.

For each cell, a one-sided binomial test evaluates whether its score direction (+ or −) remains consistent across bootstraps beyond random expectation (*H*_0_: *p* = 0.5; *H*_*a*_: *p* > 0.5). Benjamini–Hochberg correction is applied to control FDR. Cells with non-zero observed scores and FDR < 0.05 are deemed confidently assigned. Cells consistently labelled “haplotype 0-favoured” or “haplotype 1-favoured” are reported as confidently called. This bootstrap framework ensures that even if local phasing is incomplete, per-cell inference remains reliable, mitigating noise and error artifacts.

### Output

XC-ID generates a tab-separated values file containing per-cell X chromosome assignments and associated statistics. Each line includes the cell barcode, inferred active X identity ( *X*_0_, *X*_1_, or uncertain), haplotype score, binomial p-value, and Benjamini– Hochberg–adjusted p-value. The output can be directly merged with cell metadata in standard single-cell analysis frameworks, such as Scanpy or Seurat, for downstream haplotype-stratified analyses.

## Validation and Results

### Benchmark dataset and experimental setup

We validated XC-ID using single-nucleus RNA-seq data from the cerebral cortex of a female *Mus musculus* B6CASTF1/J hybrid (ENCODE accession ENCSR224RWH) with known parental phasing. Raw FASTQ reads were aligned to the GRCm39.114 genome with Y and ALT contigs removed, and variants were called using bcftools mpileup. SNPs with heterozygous genotypes and quality > 30 were retained for further processing.

After filtering, ~10000 SNPs across ~8000 cells were used as input. Simulated annealing phasing was performed using the default assign_haplotypes() function and per-cell bootstrap confidence was determined using estimate_confidence().

### Accuracy and convergence

XC-ID achieved 98% phasing accuracy relative to the ground-truth parental haplotypes (**Figure 1C**), and repeated initializations yielded nearly identical results with low standard error (**Figure S1**), confirming optimization stability of the simulated annealing algorithm. Bootstrapping filtration confidently called > 99% of cells, with > 98% of these confident cells matching the true X haplotype ratios. The predicted haplotype-ratio distribution was strongly bimodal and highly correlated with ground truth (Pearson r = 0.99; **Figure 1C**), demonstrating clear partitioning of active X identities.

### Differential expression between X lineages

We next tested whether XC-ID-partitioned X lineages capture meaningful transcriptional differences. Differential expression (DE) analysis between cells labelled as *X*_0_ and *X*_1_ was performed using Scanpy’s rank_genes_groups() Wilcoxon rank sum test (Wolf, Angerer and Theis 2018). After Bonferroni correction, five of six significant DE genes were X-linked (**Figure 1D**). The top gene, *Ogt* (O-GlcNAc transferase), is an imprinted gene that regulates the neural stem cell pool (Cheng *et al*. 2019; Andergassen *et al*. 2021; Chen *et al*. 2021). Other DE genes included the X-linked genes *Tspan7* (tetraspanin-7), *Cdr1os* (cerebellar degeneration related antigen 1), and *Las1l* (LAS1-like ribosome biogenesis factor) and the autosomal gene *Gm26871* (predicted ortholog of human miR-137 host gene), all implicated in neurodevelopmental or neurodegenerative disorders (Bassani *et al*. 2012; Xu *et al*. 2015; Hu *et al*. 2016; Wright *et al*. 2016). Another DE gene is the imprinted transcript *Crossfirre*, which maintains the inactive-X heterochromatin (Hasenbein *et al*. 2024). Together, these results show that XC-ID enables biologically coherent stratification of X chromosome lineages in scRNA-seq data.

### Runtime and scalability

We evaluated runtime across resampled datasets between 100-10000 cells and 50-10000 SNPs (**Figure 1E**). Even for the largest tested matrix (10000 cells × 10000 SNPs), the complete pipeline, including phasing, bootstrapping, and result summarization, completed in under five minutes on a standard 32-core compute node. For smaller matrices (any size with< 2000 SNPs), runtime was under 1 minute. Data loading and preprocessing were also lightweight (≈ 20 seconds for this dataset). These benchmarks highlight XC-ID’s scalability for modern cohort single-cell studies.

### Stress tests and robustness

We tested XC-ID across biologically and technically challenging scenarios, including XCI skew, escape from inactivation, reference bias, allele dropout, and extreme sparsity. XC-ID accurately assigned X chromosome identities in all test cases, given sufficient detection of confident cells (**Figure S1**).

#### XCI escape and skew

Complete and partial escape events were modelled by redistributing dominant allele reads to the minor allele. Cell assignment accuracy exceeded 98 % across all proportions of partial escape. In the complete escape scenario, accurate and confident cells were called up until 70 % of escaping SNPs, indicating robust bootstrap filtration. Cell assignment accuracy was not affected by XCI skew.

#### Transcriptomic sparsity

Downsampling SNPs, cells, and total counts showed accurate recovery of confident cells at only 14 % of SNPs, 8 % of cells, and 27 % of reads retained. Cell assignment accuracy was preserved > 99% for all scenarios with confident cell detection.

#### Technical artifacts

Simulated allele dropout and reference bias confirmed tolerance to imbalance: cell calling accuracy remained > 90% across bias below 50%. Sequencing errors (random REF/ALT flips) reduced performance gradually, with confident cells detected up to a 22% error rate.

## Discussion

XC-ID provides a lightweight framework for identifying the active X chromosome in single-cell data. Unlike genotype-dependent or heuristic methods, it reconstructs haplotypes directly from allelic expression and quantifies the confidence of each cell’s X assignment. This approach yields both speed and statistical confidence, critical for the analysis of large single-cell cohorts where sequencing quality is highly variable and mapping bias is pervasive. In benchmarking, XC-ID achieved near-perfect accuracy within minutes, requiring no prior phasing. Its robustness to XCI escape, skew, and allelic dropout makes it broadly applicable across single-cell datasets and mammalian species.

Overall, XC-ID fills a methodological gap by enabling scalable, lineage-aware single-cell studies of dosage compensation, sex-biased gene regulation, and X-linked disease mechanisms.

## Availability

XC-ID is available as a Python package with both API and command-line support. Installation instructions, documentation, and preprocessing guidelines are available at: https://github.com/jlhjiang/XC-ID.

## Acknowledgements

We thank members of the Gillis Lab at the University of Toronto and colleagues at Cold Spring Harbor Laboratory for their helpful discussions; Ahmed Abdelmoneim and Dr. Leon French for feedback on the codebase, Young June Kim for user-testing XC-ID, and Dr. Dan Levy for the conceptualization of the XC-ID algorithm.

## Author Contributions

J.L.H. Jiang (Conceptualization [supporting], Data Curation [lead], Formal Analysis [lead], Methodology [equal], Software [lead], Writing – Original Draft [lead]) and J. Gillis (Conceptualization [lead], Methodology [equal], Supervision [lead], Writing – Original Draft [supporting]).

---

**Supplementary Figure S1.**
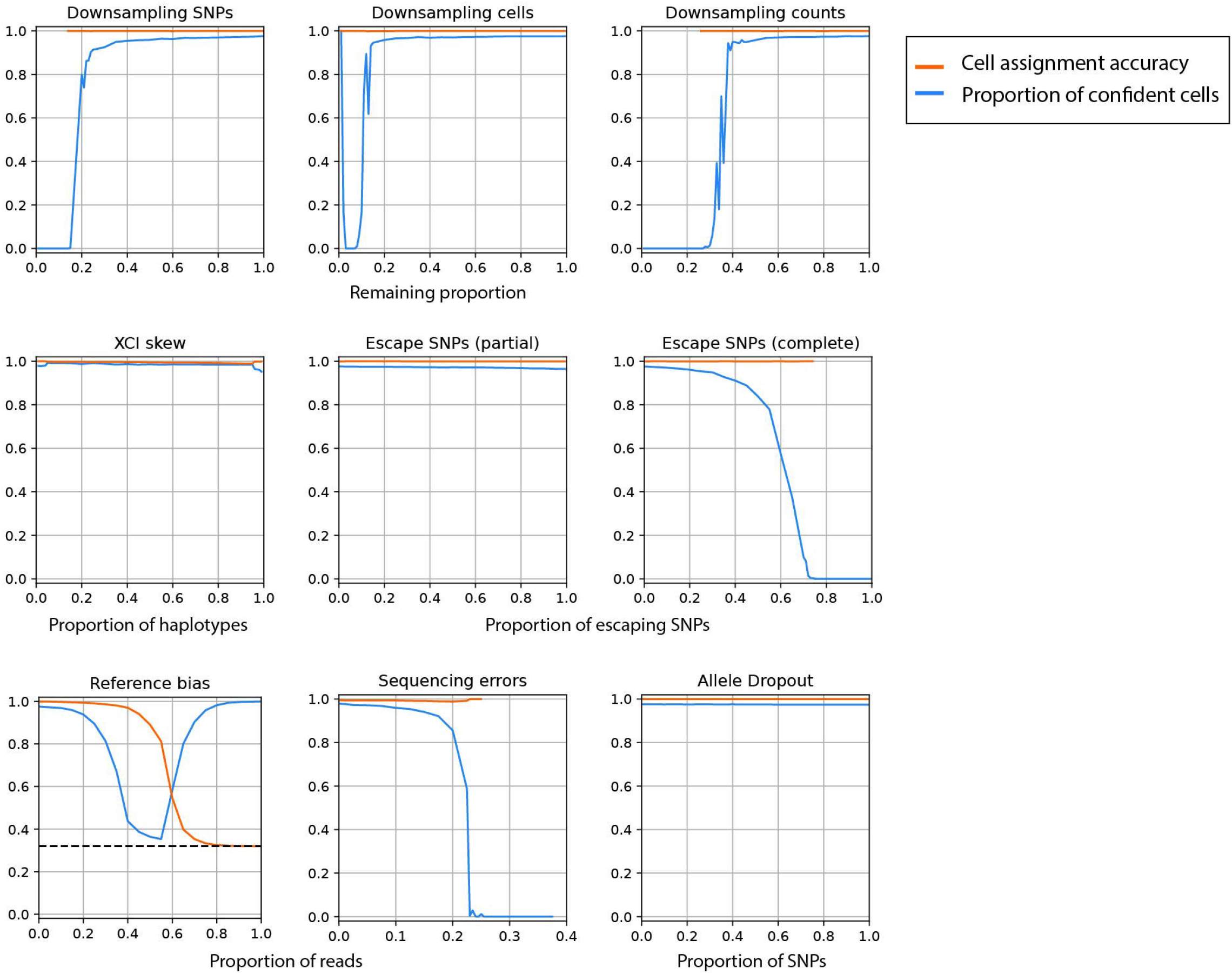
Cell-assignment accuracy (orange) and proportion of confidently called cells (blue) under simulated sparsity and technical/biological perturbations. **Top row:** effect of downsampling SNPs, cells, or total counts (sparsity) on XC-ID performance. **Middle row:** effect of XCI skew and XCI escape (biological noise). Partial escape was simulated with redistributing dominant-allele counts to the minor allele with 0.3 probability (Poisson distribution); complete escape redistributed 100% of dominant-allele counts. **Bottom row:** effect of reference-mapping bias, sequencing errors, and allele dropout (technical artifacts). In the reference-bias panel, the dashed line shows the expected REF/ALT ratio; phasing degrades toward the null as bias increases. Overall, XC-ID maintains high cell-calling accuracy provided a sufficient fraction of cells pass the confidence filter.

